# Unsupervised Spatially Embedded Deep Representation of Spatial Transcriptomics

**DOI:** 10.1101/2021.06.15.448542

**Authors:** Huazhu Fu, Hang Xu, Kelvin Chong, Mengwei Li, Kok Siong Ang, Hong Kai Lee, Jingjing Ling, Ao Chen, Ling Shao, Longqi Liu, Jinmiao Chen

## Abstract

Spatial transcriptomics enable us to dissect tissue heterogeneity and map out inter-cellular communications. Optimal integration of transcriptomics data and associated spatial information is essential towards fully exploiting the data. We present SEDR, an unsupervised spatially embedded deep representation of both transcript and spatial information. The SEDR pipeline uses a deep autoencoder to construct a low-dimensional latent representation of gene expression, which is then simultaneously embedded with the corresponding spatial information through a variational graph autoencoder. We applied SEDR on human dorsolateral prefrontal cortex data and achieved better clustering accuracy, and correctly retraced the prenatal cortex development order with trajectory analysis. We also found the SEDR representation to be eminently suited for batch integration. Applying SEDR to human breast cancer data, we discerned heterogeneous sub-regions within a visually homogenous tumor region, identifying a tumor core with pro-inflammatory microenvironment and an outer ring region enriched with tumor associated macrophages which drives an immune suppressive microenvironment.

## Introduction

Single-cell omics technologies enable measurements at single-cell resolution, and have led to discoveries of new subpopulations across various tissues, in both healthy and diseased states. However, the dissociation of tissue into single cells prior to high throughput omics data acquisition leads to cellular spatial information being lost, hindering our ability to dissect the spatial organization and intercellular interactions of individual cells. While computational tools have been developed to predict cell-cell interactions from ligand and receptor expression, they require validation using immunohistochemistry (IHC) or immunofluorescence (IF). Emerging spatial omics technologies overcome these limitations through the simultaneous measurement of gene/protein expression and spatial locations of cells. Such spatially resolved transcriptomes of histological tissues enable the reconstruction of tissue architecture and cell-cell interactions.^1,2,3,4,5,6,7,8,9^ This approach has proven valuable in many applications including studies on brain disorders,^2,10^ tumour microenvironments,^3,11^ and embryonic development.^12^

Among currently available spatial transcriptomics approaches, *in situ* capturing-based technologies such as 10x Genomics Visium and Nanostring GeoMX DSP have gained popularity owing to their accessibility and ability to profile a large number of mRNA targets within each spot. In principle, a histological section from a tissue sample is permeabilized and the released mRNA is captured by either spatially arrayed oligos on slide surfaces or by pre-hybridized RNA-target barcodes in manually defined regions of interest (ROIs). However, both technologies suffer from mRNA capture area limitations, with the smallest diameter typically being ~50μm, which is larger than a single cell. To overcome this, several computational methods have been developed to deconvolve the cell mixture of the spatial spot.^13,14,15,16,17,18,19,20^ Recently, improvements in mRNA capture methods have led to smaller subcellular capture areas that are ~1-10μm in diameter. These high-resolution spatial transcriptomics methods can obtain spatially resolved transcriptomes with increased spatial fidelity, without compromising the number of genes captured. They include Slide-seq,^8^ DBiT-seq,^9^ Stereo-seq,^5^ PIXEL-seq,^6^ and Seq-Scope,^7^ with the highest resolution (~1μm) thus far obtained by the latter three. These submicrometer-resolution methods usually require voxel binning or cell segmentation to produce a gene-by-cell expression matrix for downstream analysis. Capture area sizes have also improved and thus increased the overall cell throughput, necessitating new computational methods that can handle big spatial data.

When analyzing spatial transcriptomics data, combining both gene expression and spatial information to learn a discriminative representation for each cell or spot is crucial. However, established workflows, e.g., Seurat,^21^ still employ pipelines designed for single-cell RNA-seq analysis, which primarily focus on gene expression data and ignore the structural relationship of the spatial neighborhood. Recently, several new methods have been developed for spatial transcriptomics to overcome this limitation. For example, BayesSpace^22^ starts from a Markov random field (MRF) prior which hypothesizes that spots belonging to the same cell type should be closer to one another, and updates the model with a Bayesian approach. Giotto^23^ implements a hidden Markov random field (HMRF) model to detect domains with coherent patterns by comparing gene expression between cells and their neighbors. SpaGCN^24^ combines spatial distances and histological dissimilarities to construct a weighted graph of spots, and then integrates the graph with gene expression using a graph convolutional network (GCN) to cluster the spots. stLearn^25^ utilizes a deep learning model on the spot images to extract morphological features, on which morphological distances are calculated. It then uses the morphological distance and spatial neighborhood information to normalize the gene expression of each spot based on its identified neighbors. The normalized gene expression is then used as input for linear principal component analysis (PCA), followed by uniform manifold approximation and projection (UMAP), and spatial clustering. Notably, these methods mainly rely on PCA to extract the highly variable features of gene expression data, which involves a linear transformation, so they are unable to model complex non-linear relationships. While stLearn does utilize deep learning, it is only applied to the image modality, and the model still relies on linear PCA to extract features from the spatially normalized gene expression data. Moreover, these methods do not produce low-dimensional representations of jointly embedded gene expression and spatial information. The joint embedding of gene expression and spatial information is essential to effectively integrate both modalities for better visualization, clustering, trajectory inference, and batch integration.

In this work, we developed an unsupervised spatially embedded deep representation (SEDR) method for learning a low-dimensional latent representation of gene expression embedded with spatial information. Our SEDR model consists of two main components, a deep autoencoder network for learning a gene representation, and a variational graph autoencoder network for embedding the spatial information. These two components are optimized jointly to generate a latent representation for spatial transcriptomics data analysis. We applied SEDR on the 10x Genomics Visium spatial transcriptomics and Stereo-seq datasets and demonstrated its ability to achieve better representations for various follow-up analysis tasks, including clustering, visualization, trajectory inference, and batch effects correction.

## Results

### Overview of SEDR

SEDR learns a gene representation in a low-dimensional latent space with jointly embedded spatial information (Figure 1). Given spatial transcriptomics data, SEDR first learns a nonlinear mapping from the gene expression space to a low-dimensional feature space using a deep autoencoder network. Simultaneously, a variational graph autoencoder is utilized to aggregate the gene representation with the corresponding spatial neighborhood relationships to produce a spatial embedding. Then, the gene representation and spatial embedding are concatenated to form the final latent representation used to reconstruct the gene expression. Thereafter, an unsupervised deep clustering method^26^ is employed to enhance the compactness of the learned latent representation. This iterative deep clustering generates a form of soft clustering that assigns cluster-specific probabilities to each cell, leveraging on the inferences between cluster-specific and cell-specific representation learning. Finally, the learned latent representation can be applied towards various analysis tasks.

**Figure 1.**
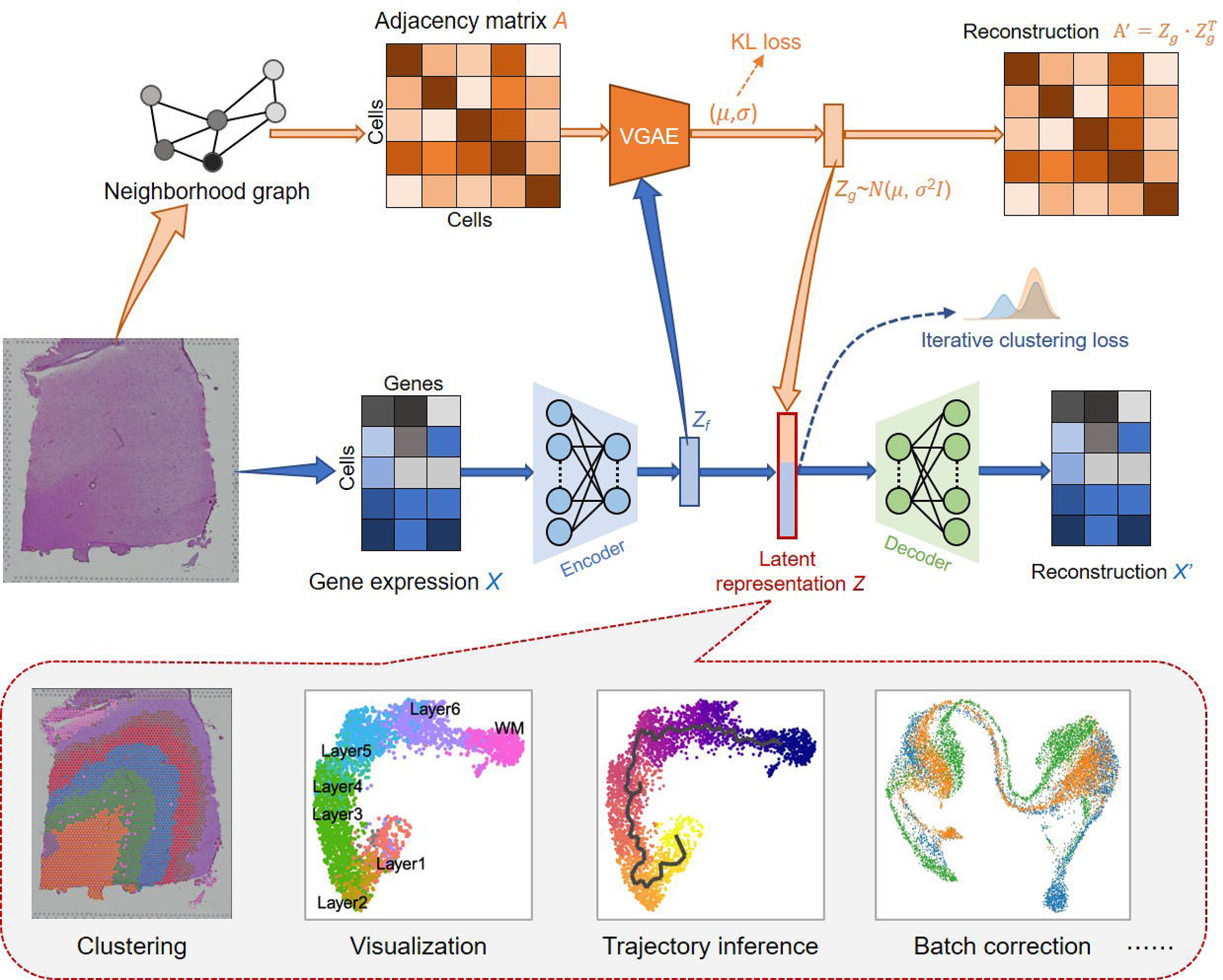
Overview of SEDR. SEDR learns a low-dimensional latent representation of gene expression embedded with spatial information by jointly training a deep autoencoder and a variational graph autoencoder. The low-dimensional embedding produced by SEDR can be used for downstream visualization, cell clustering, trajectory inference, and batch effect correction.

### Quantitative assessment of SEDR on human dorsolateral prefrontal cortex (DLPFC) dataset

To perform a quantitative comparison of SEDR with other methods, we downloaded the 10x Genomics Visium spatial transcriptomics data and the manually annotated layers for LIBD human dorsolateral prefrontal cortex (DLPFC) data.^2^ The LIBD data includes 12 slices from the human DLPFC, which span six cortical layers plus white matter. We chose this dataset because the human DLPFC has clear and established morphological boundaries which can serve as the ground truth. We first applied the standard Seurat pipeline^21^ to process and cluster cells using only expression profiles and set the result as the benchmarking baseline to investigate the extent to which spatial information improves cell clustering. As Giotto,^23^ stLearn,^25^ SpaGCN,^24^ and BayesSpace^22^ integrate spatial information and RNA-seq data for clustering, we also applied them with recommended default parameters to the same dataset for benchmarking against SEDR.

In brain slice 151673 (Figure 2A) with 3,639 spots and 33,538 genes, SEDR and BayesSpace achieved the best performance in terms of both layer borders and adjusted rand index (ARI). When comparing the results on all 12 DLPFC samples, SEDR had the second highest mean ARI (0.427) (Figure 2A, bottom right), but the difference between SEDR and the top performer BayesSpace (0.457) was not significant (Mann-Whitney U Test:^27^ p-value=0.55). It should be noted that BayesSpace’s clustering algorithm is optimized for spatial omics, while SEDR is a dimension reduction method with its objective being to find the best latent representation. SEDR followed by Leiden clustering, which was not specifically designed or optimized for clustering spatial omics, achieved comparable clustering performance to BayesSpace. This indicated that SEDR latent representations effectively integrate gene expressions and spatial information for capturing inter-cluster differences. Coupling SEDR with clustering algorithms that are better-suited for spatial omics can be expected to further improve the clustering accuracy. Furthermore, in contrast to BayesSpace, which does not produce latent representations, SEDR-derived embeddings can be used for not only clustering but also various downstream analysis tasks such as UMAP visualization, trajectory inference, and batch effect correction, thus providing more flexibility and utility. Similar to SEDR, SpaGCN also uses a GCN to process spatial transcriptomics data. Moreover, it incorporates histological information, which is not included in SEDR. However, the clustering performance of SEDR is better than that of SpaGCN (Mann-Whitney U Test, p-value < 0.05). stLearn also integrates histological data, but the performance is likewise poorer. This may indicate that the current approaches utilized by SpaGCN and stLearn to incorporate histological data are not optimal. To make full use of the histological information, we may need to treat it as a separate data modality and use dedicated multi-view algorithms for integration.

**Figure 2.**
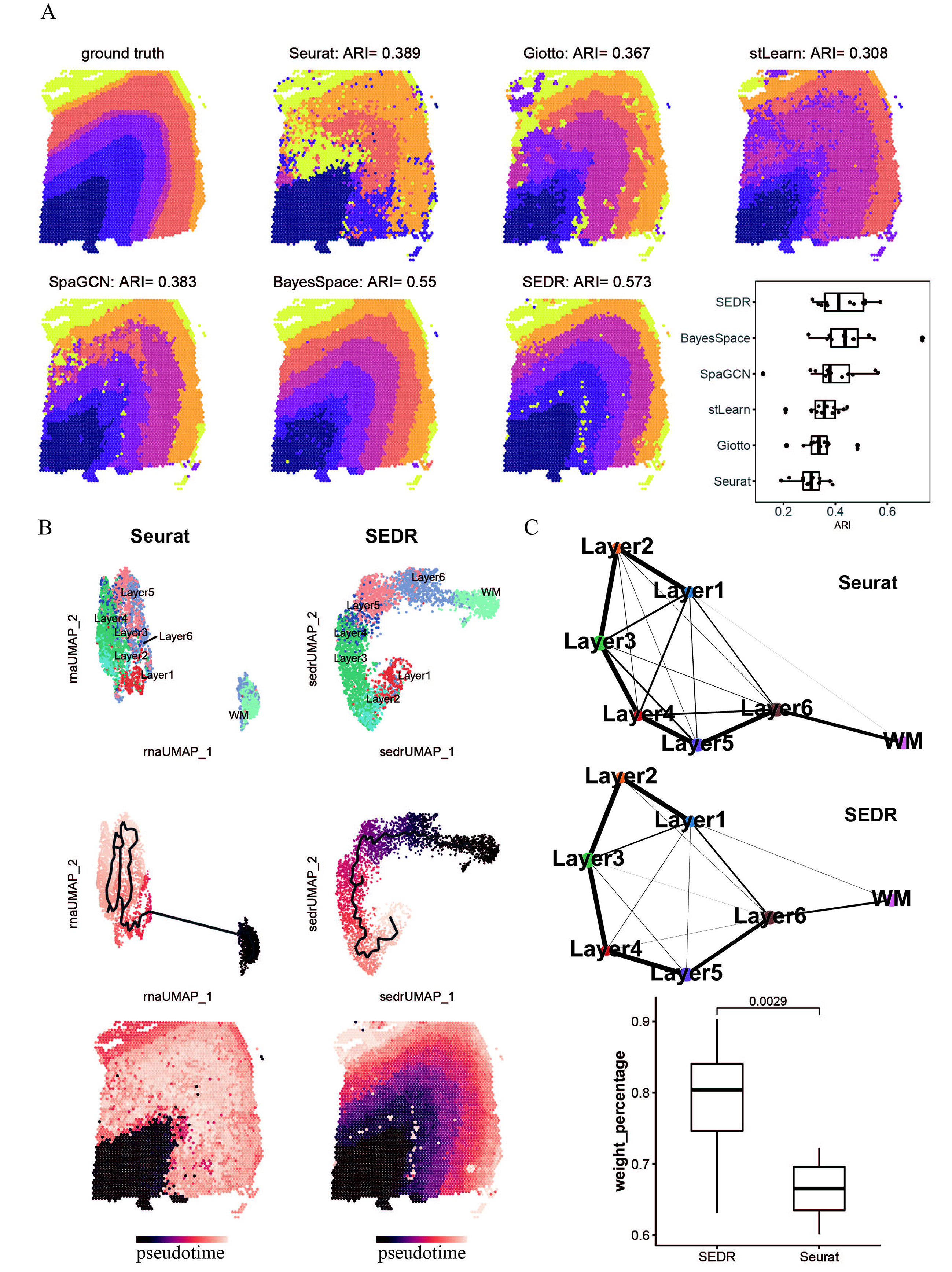
Quantitative assessment of SEDR on the human dorsolateral prefrontal cortex (DLPFC) dataset. A) Ground-truth segmentation of cortical layers; clustering results of Seurat, Giotto, stLearn, SpaGCN, BayesSpace, and SEDR on DLPFC slice #151673; and adjusted rand index (ARI) of various cluster sets on the 12 DLPFC slices. B) UMAP visualization and Monocle3 trajectory generated using the Seurat-derived PCA embedding (left) and SEDR embedding (right). Monocle pseudotimes visualized on UMAP plots (middle) and spatial co-ordinates (bottom). C) PAGA graphs generated using the Seurat-derived PCA embedding (top) and SEDR embedding (middle). The ratios of the sum of weights of correctly inferred PAGA edges to the sum of weights of all edges produced by SEDR and Seurat (bottom).

SEDR generates a set of low-dimensional representation features which can be used in various downstream analyses, such as trajectory inference.^28^ In our experiments, we used Monocle3^29^ to perform trajectory inference on sample 151673 with the Seurat output (RNA-only) and the low-dimensional SEDR representation features. We found that SEDR showed significantly improved performance over Seurat (Figure 2B). In the UMAP plot of SEDR’s output, cells belonging to different layers were well-organized, and when we selected white matter (WM) as the root, the pseudo-time reflected the correct “inside-out” developmental ordering of cortical layers (Figure 2B). This demonstrated that, compared to RNA-only analyses, incorporating spatial information enabled SEDR to generate a better latent representation summarizing the spatial transcriptomics data. We further confirmed our observations with another trajectory inference method named partition-based graph abstraction (PAGA),^30^ using the SEDR-derived latent space embedding instead of UMAP coordinates (Figure 2C). The PAGA results showed that adjacent cortical layers tend to share greater similarity, suggesting that spatial adjacency is linked with transcriptomic and even functional similarity. Notably, the trajectory was concordant with the chronological order of cortex development.^31,32,33^ We then compared the PAGA graphs generated using the Seurat-derived principal components and SEDR embeddings. For each of the 12 DLPFC slices, we calculated the ratio of the edge weights between adjacent cortical layers to the total sum of the weights of all edges. We found a significantly higher ratio for SEDR compared to Seurat (Mann-Whitney U test p-value < 0.05) (Figure 2C, bottom).

### SEDR corrects for batch effects

The proliferation of spatial omics applications is generating ever increasing volumes of spatially resolved omics data across different labs. However, differences in protocols and technologies complicate comparisons and data integration when trying to achieve consensus on spatially resolved tissue atlases. As with single-cell RNA-seq (scRNA-Seq), removing batch effects in spatial omics datasets is a significant challenge. To date, no methods are available for this. Here, we demonstrate that SEDR can learn joint embeddings across multiple batches and project them into a shared latent space. Furthermore, SEDR employs a deep embedded clustering (DEC) loss function that enables it to retain biological variations while reducing technical variations. We evaluated the batch correcting performance of SEDR on the DLPFC datasets. We first assessed the batch variations among the twelve datasets and selected three sets (151507, 151672, 151673) which exhibited substantial batch effects. The common cortical layers from different batches were separated, as shown in the UMAP plot (Figure 3A). We first applied Harmony to remove the batch effects due to its superior performance in scRNA-seq data integration.^34^ Harmony was able to mix batches while keeping different layers apart. However, when we zoomed into the individual layers, we found distinct batch-specific subclusters, suggesting that the batch effects were not completely removed (Figure 3B). Next, we tested SEDR and found that the batch effects were substantially reduced (Figure 3C). Common layers across batches were brought very close and were well-aligned, while different layers were minimally mixed. Further application of Harmony on the SEDR embeddings evenly mixed the batches while maintaining separation between layers (Figure 3D). Notably, batch-specific clusters were no longer present within individual layers. This showed that the combination of SEDR with Harmony effectively removed the batch effects. Among the other spatial omics analysis methods, only stLearn is able to produce a latent space embedding which can be fed to Harmony for batch correction. Therefore, we benchmarked SEDR against stLearn. As stLearn is unable to jointly project different batches to a shared latent space due to its requirement of histological images as input, we generated latent space embeddings from each dataset and then concatenated them for Harmony integration. The results showed that batches were not well mixed and the layers were poorly separated (Figure 3E). In conclusion, SEDR combined with Harmony outperforms both Harmony alone and stLearn with Harmony, and can serve as an effective method for batch correction of spatial omics data.

**Figure 3.**
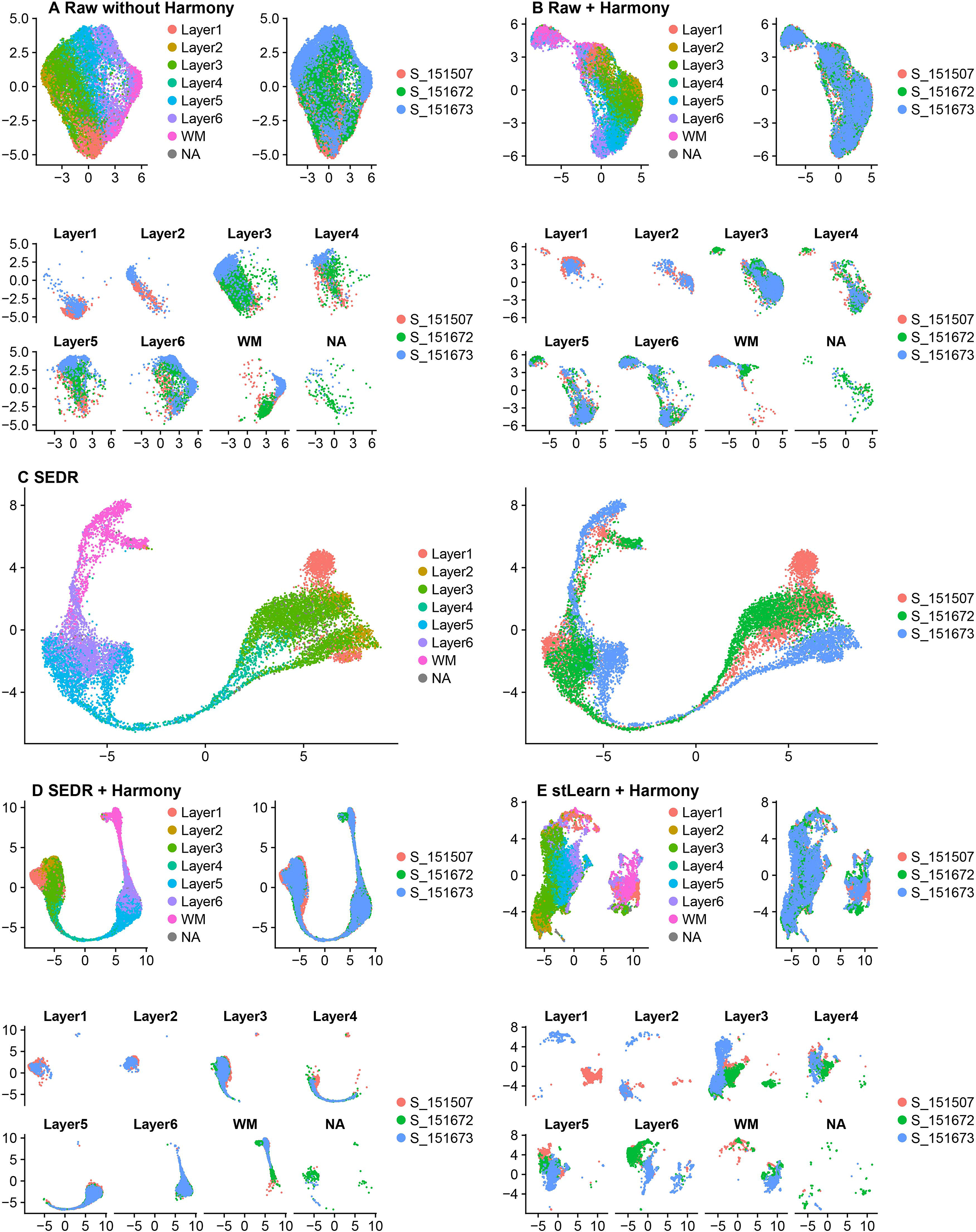
Batch effects present in DLPFC dataset and assessment of SEDR’s performance on batch correction. A) Slices #151507, #151672 and #151673 showed substantial inter-slice variations before batch effect correction. UMAP plots colored by ground-truth cortical layers (left), slices (right), split by layers and colored by slices (bottom). B) Harmony alone was unable to remove the batch effects present. C) SEDR alone substantially reduced the batch effects. D) SEDR combined with Harmony effectively corrected the batch effects. E) stLearn combined with Harmony was unable to correct the batch effects.

### Dissecting tumor heterogeneity and immune microenvironments using SEDR

Intratumoral heterogeneity in cancer complicates effective treatment formulations and is associated with poor survival prospects.^35^ Spatial transcriptomics is an effective tool for dissecting and characterizing intratumoral heterogeneity and tumor-immune crosstalk. We tested SEDR on the 10x Visium spatial transcriptomics data for human breast cancer, which is known for its high intratumoral and intertumoral differences. To aid in the interpretation of SEDR results, we performed manual pathology labeling based on H&E staining. It should be noted that, unlike the cerebral cortex which has clear and established morphological boundaries, tumor tissues are highly heterogeneous and encompass complex microenvironments. Manual labeling solely based on tumor morphology is inadequate for characterizing such complexity. Based on pathological features, we manually segmented the histological image into twenty regions, which we then grouped into four main morphotypes: ductal carcinoma *in situ*/lobular carcinoma *in situ* (DCIS/LCIS), healthy tissue (Healthy), invasive ductal carcinoma (IDC), and tumor surrounding regions with low features of malignancy (Tumor edge) (Figure 4A top left, Supplementary Figure 1A). Visually, all five methods agreed with the manual annotations at the macroscopic level (Figure 4A). Nevertheless, the SEDR clusters presented a smoother segmentation compared to other methods, while those derived by Seurat, stLearn, and SpaGCN appeared fragmented with irregular boundaries. Notably, SEDR found more sub-clusters within the tumor regions, while other methods were prone to dividing the healthy regions into subclusters, given that all methods were set to generate the same number of clusters. Specifically, within the seemingly homogenous tumor region DCIS/LCIS_3, SEDR separated an outer “ring” (Figure 4A, SEDR cluster 7) from the tumor core (Figure 4A, SEDR cluster 3). These SEDR clusters indicated transcriptionally and spatially distinct compartments within the visually homogenous tumor regions. In addition to clustering analysis, we also employed the Seurat3 ‘anchor’-based integration workflow to perform probabilistic transfer of annotations from scRNA-seq reference data for human breast cancer^36^ to the spatial data. For each spot, we obtained a probabilistic classification for each of the scRNA-seq derived classes (Figure 4B, Supplementary Figure 1B). The transferred class probabilities were able to delineate the tumor regions and regions where immune cells or fibroblasts were present, which were useful for further dissecting the tumor microenvironment.

**Figure 4.**
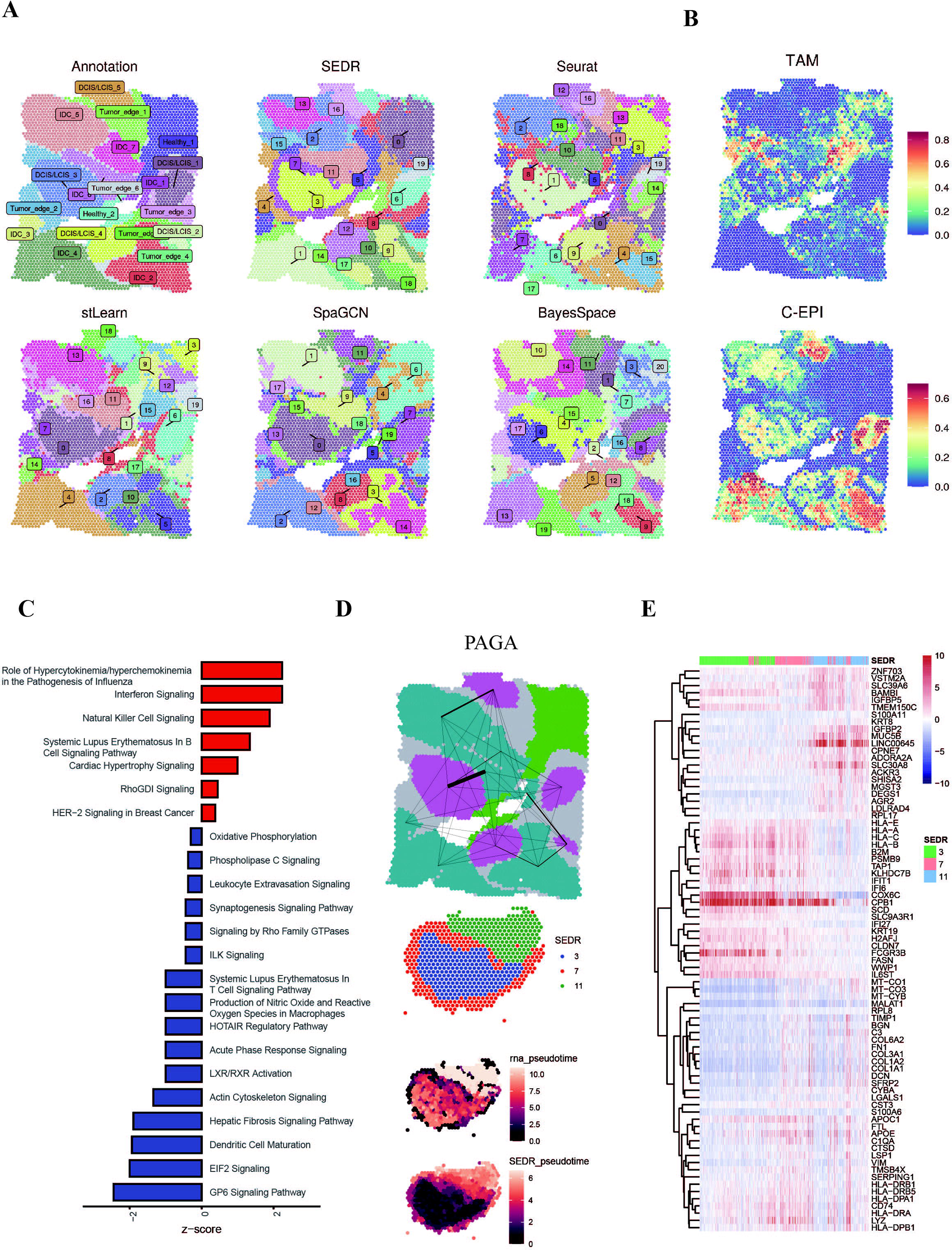
Application of SEDR on 10x Visium spatial transcriptomics data of human breast cancer. A) Manual pathology labeling based on H&E staining (annotation); clustering results of SEDR, Seurat, stLearn, SpaGCN, and BayesSpace. B) Seurat3 ‘anchor’-based integration workflow was used to perform probabilistic transfer of annotations from a reference scRNA-seq data of human breast cancer to the spatial data. This gives a probabilistic classification of the scRNA-seq derived classes for each spot. The probabilities of tumor associated macrophage (TAMs) and cycling epithelials (C-EPI) were visualized. C) Pathways enriched by genes differentially expressed between SEDR clusters 3 (core) and 7 (outer ring). Red bars represent pathways upregulated in cluster 3. D) Trajectory analysis results using PAGA (top) and Monocle3 (bottom). The PAGA graph predictions of the inter-relatedness between the manually annotated DCIS/LCIS and IDC regions. The edge width denotes connectivity strength, thus indicating the likelihood of an actual connection being present. Monocle3 inferred the pseudo-times of spots in SEDR clusters 3, 7, and 11 using the Seurat-derived PCA embedding (termed “rna_pseudotime”) and SEDR embedding (termed “SEDR_pseudotime”). E) Heatmap of genes with expression changes along the Monocle-derived pseudo-time.

To further characterize the transcriptional differences between SEDR cluster 3 (tumor core) and cluster 7 (tumor edge) of DCIS/LCIS_3 region, we performed differential expression analysis followed by pathway enrichment analysis (Figure 4C). In cluster 3, we observed the upregulation of interferon signaling pathways (IFIT1, IFITM1, IFITM3 and TAP1) and NK or neutrophil activities (FCGR3B and TNFSF10) (Figure 4C, Supplementary Figure 2E). In addition, RHOB was upregulated in this region, pointing towards reduced metastatic potential.^40^ Cluster 3 represented a region where cancer growth was limited by pro-inflammatory immune responses. On the other hand, in cluster 7, we observed the presence of TAMs (Figure 4B, Supplementary Figure 2D), memory B cells (IGHG1, IGHG3, IGHG4, IGLC2 and IGLC3) and fibroblasts (COL1A1, COL1A2, COL3A1, COL5A1, COL6A1, COL6A2 and FN1) (Figure 4C, Supplementary Figure 2E). Upregulated cathepsin activity (CTSB, CTSD and CTSZ) and complement pathway (C1QA, C1S) indicated protumor activity by the TAMs in this region.^41,42,43^ Upregulation of actin cytoskeleton signalling also suggested higher metastasis potential of cluster 7 (Figure 4C). Moreover, upregulated cathepsin activity and metalloproteinase inhibitors (TIMP1 and TIMP3) also indicated disturbance in the extracellular matrix integrity (Supplementary Figure 2E). Overall, cluster 7 represented a region with an immune-suppressed pro-tumor microenvironment and high potential for cancer metastasis.

A number of driving forces have been hypothesized as responsible for the metastatic transition of tumor cells from a pre-invasive state to invasive carcinoma, including pro-tumor immune microenvironments and reduced cell-cell interactions within the tumor.^37^ In this study, we employed PAGA to infer the inter-relatedness between the manually annotated tumor regions to trace the metastatic transition process. The PAGA graph generated using the SEDR embeddings suggested that DCIS_LCIS_3 was related to the neighboring IDC_6 region (Figure 4D). The differentially expressed genes (DEGs) and enriched pathways of DCIS_LCIS_3 compared to all other DCIS_LCIS regions showed that DCIS_LCIS_3 had more immune infiltrates (Supplementary Figure 2A, 2B, 2C), particularly tumor associated macrophages (TAMs) (Figure 4B, top), while the other DCIS_LCIS regions were mainly comprised of actively dividing/cycling epithelial cells (Figure 4B, bottom) with upregulated glycolytic and metabolic processes (Supplementary Figure 2C). TAM infiltration is known to be strongly associated with poor survival rate in solid tumor patients due to its promotion of tumor angiogenesis and induction of tumor migration, invasion and metastasis.^38,39^ We thus performed Monocle3 analysis to infer the pseudo-time of the transition from DCIS_LCIS_3 to IDC_6. As DCIS_LCIS_3 and IDC_6 coincided with SEDR clusters 3, 7, and 11 (Figure 4A, 4D), we applied Monocle3 on these three clusters and set cluster 3 (tumor core) as the starting point (Figure 4D bottom). Monocle3 analysis showed that pseudo-time derived from SEDR embeddings better traced the inside-out tumor progression compared to that from Seurat PCA embeddings. We subsequently identified genes that changed expression along the Monocle3 pseudo-time and revealed sequential waves of gene regulation along the trajectory (Figure 4E).

In summary, SEDR analysis revealed intratumoral heterogeneity within visually homogenous tumor regions and revealed the tumor outer ring (cluster 7) with TAM infiltration and cancer associated fibroblasts (CAFs), both of which have been reported to facilitate tumor spread.^44,45^ SEDR also enabled the mapping of a molecular trajectory from the tumor core to its neighboring invasive region, demonstrating the transition from a pro-inflammatory to an immune-suppressive microenvironment, which may contribute to tumor metastasis.

### SEDR can handle spatial transcriptomics of high resolution

Currently available spatial omics technologies, including 10x Visium Spatial Omics, Nanostring GeoMX DSP, SLIDE-seq^4^, and DBIT-seq^46^, do not provide single-cell resolution, with each capture spot containing 1 to 10 cells. However, newly emerging methods such as Stereo-seq^5^, PIXEL-Seq^6^, and Seq-Scope^7^ can achieve submicrometer and thus subcellular resolution. With continued technology advancement, the spatial resolution and number of cells detected per tissue will significantly improve, producing large datasets with high throughput. As such, we evaluated SEDR’s performance on one type of such data produced by Stereo-seq from mouse olfactory bulb tissues (Figure 5). The coronal section of a mouse olfactory bulb contains the olfactory nerve layer (ONL), glomerular layer (GL), external plexiform layer (EPL), mitral cell layer (MCL), internal plexiform layer (IPL), granule cell layer (GCL), and rostral migratory stream (RMS) (Figure 5A). We performed unsupervised clustering using the Seurat-derived principal components and SEDR-derived embeddings to computationally reconstruct the spatial identity of the olfactory bulb tissues. Compared to Seurat clusters, those produced by SEDR better reflected tissue organization and were more consistent with known anatomical layers (Figure 5B, 5C). We also performed quantitative assessment using local inverse Simpson’s index (LISI) and found that SEDR produced significantly lower LISI than Seurat, implying better spatial separation by SEDR (Figure 5D).

**Figure 5.**
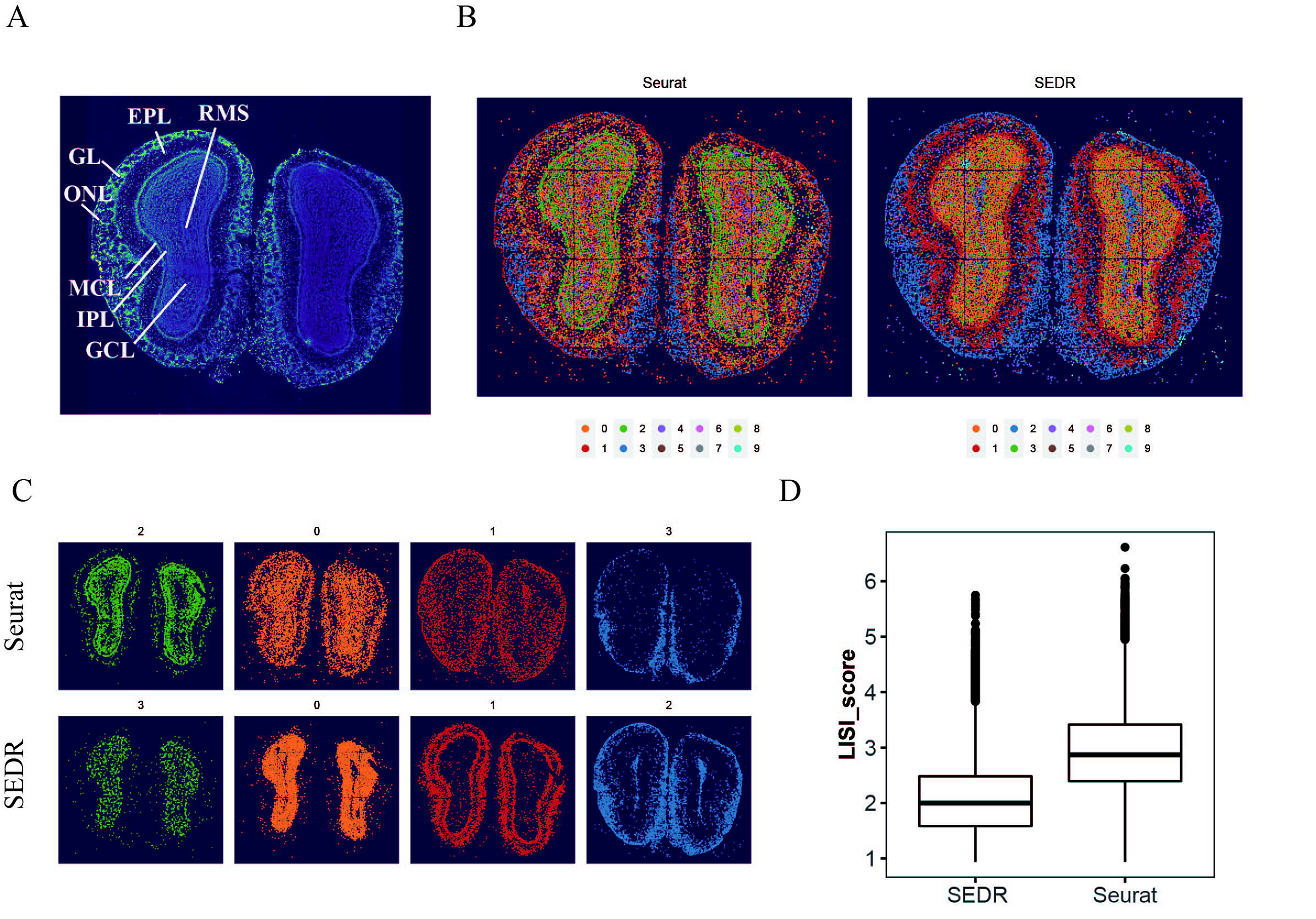
Application of SEDR on Stereo-seq spatial transcriptomics data of mouse olfactory bulb tissue sections. A) Laminar organization of DAPI-stained mouse olfactory bulb. B) Unsupervised clustering of the spatial voxels analyzed by Seurat and SEDR. C) The four clusters with the highest numbers of voxels were selected and visualized. D) Quantitative comparison of Seurat and SEDR clusters using local inverse Simpson’s index (LISI).

## Discussion

Cell type heterogeneity is a feature of both healthy and diseased tissue. Capturing this heterogeneity, coupled with its spatial arrangement in the tissue, is crucial when studying the roles of cells and their cross-talk. Spatial omics technologies represent the state-of-the-art approaches for capturing omics data with corresponding spatial information from tissue samples. In this paper, we have introduced SEDR, which leverages on cutting edge graph neural network techniques to achieve a better representation of spatial omics data that can be used for clustering and further downstream analyses. SEDR first learns a low-dimensional latent space representation of the transcriptome information with a deep autoencoder network, which is then aggregated with spatial neighborhood information by a variational graph autoencoder to create a spatial embedding. This spatial embedding is then concatenated with the gene expression to be decoded to reconstruct the final gene expression for further analyses. We first demonstrated the efficacy of SEDR in delineating the different cerebral cortex layers with higher clarity than competing methods, and recapitulated the associated development order by using the joint latent representation with Monocle3.

To enhance the analytical power and resolution of spatial omics, we need to integrate multiple datasets from the same tissue. Similar to single-cell transcriptomic data, spatial omics datasets generated in different batches also contain batch-specific systematic variations that present a challenge to batch-effect removal and data integration. In our study, we demonstrated that by combining SEDR and Harmony, we were able to effectively remove batch effects present. In the future, we will integrate Harmony into the SEDR workflow.

Spatial omics technologies such as Stero-seq are able to measure a large number of cells in a single experiment through high spatial resolutions and large tissue sizes. In the near future, we expect to see ever-increasing throughput from spatial omics experiments, which will result in spatial omics big data that will pose significant challenges to data analysis and integration. Computational methods that employ GCNs require the entire graph to be loaded into GPU memory, which inhibits their application to very large datasets. We will improve the memory efficiency of SEDR using a GCN mini-batch or parallel techniques to construct large-scale graphs for spatial omics data of high throughput and resolution. Furthermore, technologies with a capture spot size smaller than the diameter of a cell will also require new computational methods that can accurately delineate cells based on capture spots. In the future, we will integrate cell segmentation based on H&E or DAPI staining into the SEDR workflow.

The current SEDR methodology focuses on gene expression and spatial information, and does not make use of histological images. Contemporary methods such as SpaGCN and stLearn use histological images as input, but in a suboptimal fashion, as demonstrated in our study. Specifically, SpaGCN utilizes histological image pixels as features by calculating the mean color values from the RGB channels directly. However, the pixel values are easily affected by noise and cannot provide semantic features for cell analysis. A more effective approach can be to adopt a deep CNN model which can learn high-level representations for histological images. stLearn introduces a deep learning model to extract image features of the spots and integrates them with the spatial location and gene expression. However, stLearn employs a model pre-trained on natural images, and does not fine-tune the network for histological images. In the future, we will incorporate histological images as an additional modality into the SEDR model. We will add an image autoencoder network to learn image features, and jointly learn the latent representation by integrating gene expression, image morphology, and spatial information.

In summary, SEDR is a promising new approach that builds an integrated representation of cells using both transcriptomic data and spatial coordinates. SEDR-derived low-dimensional embedding enables more accurate clustering, trajectory inference and batch effect correction. Our model is also able to handle spatial transcriptomics with capture spot sizes ranging from 50μm to less than 1μm. Furthermore, we applied SEDR on human breast cancer to reveal heterogeneous sub-regions within the seemly homogenous tumor region and shed light on the role of immune microenvironments on tumor invasiveness.

## Methods

### Dataset preprocessing

Our SEDR method takes spatial transcriptomic gene expressions and spatial coordinates as inputs. The raw gene expression counts are first normalized using the respective library sizes (by normalize_total in Scanpy (v.1.5.0)), with very highly expressed genes excluded when computing the normalization factor (size factor) for each cell^47^. PCA is then applied to extract the first 200 principal components to generate the initial gene expression matrix.

### Graph construction for spatial transcriptomics data

To create a graph representing the cell–cell spatial relationships in spatial transcriptomics data, we calculate the Euclidean distances between cells using the image coordinates, and select the top 10 nearest neighbors of each cell to construct an adjacency matrix. The adjacency matrix, denoted by *A*, is a symmetric matrix, where *A*_*ij*_ = *A*_*ji*_ = 1 if *i*, and *j* are neighbors, and 0 otherwise.

### Deep autoencoder for latent representation learning

The latent representation of gene expression is learned using a deep autoencoder. The encoder part, consisting of two fully connected stacked layers, generates a low-dimensional representation *Z*_*f*_ ∈ ℝ^*N* × *D*_*f*_^ from the input gene expression matrix *X* ∈ ℝ^*N*×*M*^. Meanwhile, the decoder part with one fully connected layer reconstructs the expression matrix *X*′ ∈ ℝ^*N*×*M*^ from the latent representation *Z* ∈ ℝ^*N*×*D*^, which is obtained by concatenating the low-dimensional representation *Z*_*f*_ and spatial embedding *Z*_*g*_ ∈ ℝ^*M*×*D*_*g*_^, where *N* is the number of cells, *M* is the number of input genes, and *D*_*f*_, *D*_*g*_, *D* are the dimensions of the low-dimensional expression representation learned by the encoder, the spatial embedding learned by the GCN, and the final latent representation of SEDR, respectively with *D* = *D*_*f*_ + *D*_*g*_. The objective function of the deep autoencoder maximizes the similarity between the input gene and reconstructed expressions measured by the mean squared error (MSE) loss function Σ(*X* – *X*′)^2^.

### Variational graph autoencoder for spatial embedding

SEDR utilizes a variational graph autoencoder^48^ (VGAE) to embed the spatial information of neighboring cells. With the adjacency matrix *A*, and its degree matrix *D*, the VGAE learns a graph embedding *Z*_*g*_ with the following format: *g*:(*A*, *Z*_*f*_)→*Z*_*g*_ where *Z*_*f*_ is the node/gene representation from the deep autoencoder. The inference part of the VGAE is parameterized by a two-layer GCN^49^ :

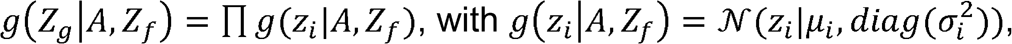

where *μ* = *GCN*_*μ*_(*A*,*Z*_*f*_) is the matrix of mean vectors, and *logσ* = *GCN*_*σ*_(*A*, *Z*_*f*_). The two-layer GCN is defined as:

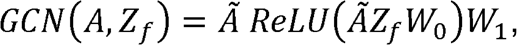

with a weight *W*_*i*_ and symmetrically normalized adjacency matrix 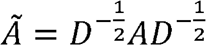. The spatial embedding *Z*_*g*_ and reconstructed adjacency matrix *A′* are generated as:

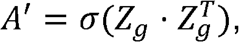

with *Z*_*g*_ = *GCN* (*A*,*Z*_*g*_). The objective of the VGAE is to minimize the cross-entropy (CE) loss between the input adjacency matrix *A* and reconstructed adjacency matrix *A′* while simultaneously minimizing the Kullback-Leibler (KL) divergence between (*g*(*Z*_*g*_│*A*,*Z*_*f*_)) and the Gaussian prior:

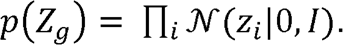

### Batch effect correction for spatial transcriptomics

Spatial relationships only exist within single spatial transcriptomic measurement; cells/spots from different transcriptomic measurements have no direct spatial relation. Let *A*^*k*^ and *Z*^*k*^_*f*_ denote the adjacency matrix and deep gene representation of spatial omics *k*, we then create a block-diagonal adjacency matrix *A*_*k*_ and concatenate the deep gene representation in the cell dimension, as:

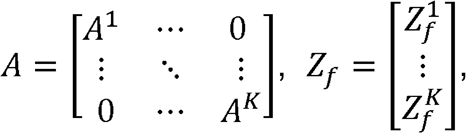

where *K* is the number of spatial omics. Based on this formulation, we transform different spatial omics datasets (of potentially different sizes) into multiple graph instances in the form of one block-diagonal adjacency matrices as inputs to SEDR.

To remove batch effects and enhance the compactness of its latent representation, SEDR employs an unsupervised deep embedded clustering (DEC) method^26^ to iteratively group the cells into different clusters. To initialize the cluster centers, we employ the KMeans of scikit-learn on the learned latent representations. The number of clusters is pre-defined as a hyperparameter. With the initialization, DEC improves the clustering using an unsupervised iterative method of two steps. In the first step, a soft assignment *q*_*ij*_ of latent point *z*_*i*_ to cluster center *μ*_*j*_ is calculated using the Student’s t-distribution:

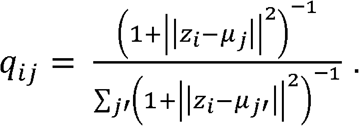

In the second step, we iteratively refine the clusters by learning from their high confidence assignments with the help of an auxiliary target distribution *p* based on *q*_*ij*_:

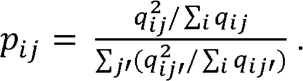

Based on the soft assignment *q*_*ij*_ and auxiliary target distribution *p*_*ij*_ an objective function is defined using the KL divergence:

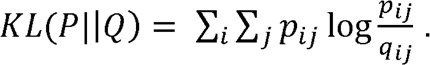

The SEDR parameters and cluster centers are then simultaneously optimized by using stochastic gradient descent (SGD) with momentum.

### Seurat

Raw mRNA counts were preprocessed to remove low-quality genes and sctransformed to remove technical artifacts and normalize the data.^50^ We then ran PCA to extract the top 30 principal components (PCs) and used them to calculate the shared nearest neighbors (SNNs). Then, the Louvain clustering algorithm was used to identify clusters with the SNN networks. We tried clustering at different resolutions to obtain the same number of clusters as the ground truth layers.

### SpaGCN, stLearn, BayesSpace, Giotto

We ran these methods with the recommended parameters and set each one to generate the same number of clusters as the ground truth layers. The stLearn-derived low-dimensional embedding was used for downstream UMAP visualization and Harmony batch correction.

### Evaluation metrics for clustering

For datasets with cell-type labels (e.g., DLPFC), we employed ARI to compare the performance of different clustering algorithms. ARI calculates the similarity between the clustering labels predicted by the algorithm and reference cluster labels as:

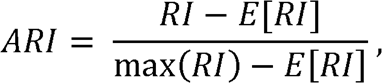

where the unadjusted rand index (RI) is defined as *RI* = (*a* + *b*)/*C*^2^_*n*_ with *a* being the number of pairs correctly labeled as coming from the same set, being the number of pairs correctly labeled as not in the same set, and *C*^2^_*n*_ being the total number of possible pairs. *E[RI]* is the expected *RI* of random labeling. A higher ARI score indicates better performance.

### Monocle3

On the DLPFC #151673 slice and breast cancer data, we ran Monocle3 using both the Seurat and SEDR outputs. For Seurat, we ran the standard pipeline to obtain the UMAP. For SEDR, we first extracted the low-dimensional embedding and then used the uwot package to calculate the UMAP. We then ran Monocle3 on both UMAPs using the recommended parameters and set white matter (WM) as the starting point to generate the pseudo-time. Finally, we used the Moran_I test to detect significant genes that showed correlations with the pseudo-time.

### Leiden clustering, PAGA trajectory, and UMAPs

We used the Scanpy (v.1.5.0) package to compute the Leiden clustering, partition-based graph abstraction (PAGA), and uniform manifold approximation and projection (UMAP) from SEDR-derived joint embeddings of gene expression and spatial information. Briefly, we used SEDR embeddings to compute neighborhood graphs with 15 as the number of neighbors and Euclidean distance as the distance measure. To obtain the same number of unique Leiden clusters, grid-searching on the Leiden clustering resolutions between 0.2 and 2.5 was performed at intervals of 0.05/0.01. Subsequently, PAGA was applied to quantify the connectivity between Leiden clusters. Finally, the cluster positions suggested by PAGA were used to compute the UMAP for visualization.

### Harmony

Harmony was used to correct batch effects on low-dimensional embeddings. For SEDR, we used latent space embeddings as input. For the raw data and stLearn, we used the PCA embeddings. We treated different samples as different batches, and set all other parameters to their default values. For each method, the uncorrected embeddings and batch-corrected Harmony embeddings were used for UMAP visualization and analysis.

### Prediction of cell type composition of 10x Visium spatial spot

We downloaded a published scRNA-seq dataset of human breast cancer^36^ as reference, and ran Seurat to find transfer anchors between the reference and our Visium spatial data. Cell types in the reference were then assigned to the spatial spots by label transferring. We removed cell types with probabilities equal to 0 for all spots.

### Differential expression analysis and pathway analyses

We used Seurat to identify DEGs. Genes with adjusted p-values < 0.05 were used as the input for QIANGEN Ingenuity Pathway Analysis (IPA). For IPA results, pathways with positive or negative z-scores were plotted.

### Raw data processing of Stereo-seq data

Fastq files were generated using the MGI DNBSEQ-Tx sequencer. Coordinate identities (CIDs) and unique molecular identifiers (UMIs) were encoded in the forward reads (CID: 1-25bp, UMI: 26-35bp), while the reverse reads consisted of the cDNA sequences. CID sequences in the forward reads were first mapped to the designed coordinates of the *in situ* captured chip, allowing one base mismatch to correct for sequencing and PCR errors. Reads with UMIs containing either N bases or more than two bases with quality scores lower than 10 were filtered out. The CIDs and UMIs associated with each read were appended to each read header. Retained reads were then aligned to the reference genome (mm10) using STAR^51^, and mapped reads with MAPQ ≥10 were counted and annotated using an *in-house* script (available at https://github.com/BGIResearch/handleBam). UMIs with the same CIDs and gene loci were collapsed together, allowing for one mismatch to correct for sequencing and PCR errors, to give the final gene expression matrix.

### Local inverse Simpson’s index (LISI)

We first used Seurat and SEDR to generate cell clusters for the stereo-seq data, and then the R “lisi” package to calculate the LISIs using spatial coordinates as X and the clustering results of Seurat and SEDR as meta data.

## Data availability

(1) LIBD human dorsolateral prefrontal cortex (DLPFC) Data (http://spatial.libd.org/spatialLIBD/); (2) 10x visium spatial transcriptomics data of human breast cancer and Stereo-seq of mouse olfactory bulb are at https://github.com/JinmiaoChenLab/SEDR/ (3) Analysis results and scripts to reproduce the results are at https://github.com/JinmiaoChenLab/SEDR/

## Software availability

SEDR was written in Python using the PyTorch library. An open-source implementation of SEDR has been released on https://github.com/HzFu/SEDR

## Acknowledgements

This research was supported by funding from Singapore Immunology Network (SIgN), A * STAR, Singapore.

## Author contributions

Huazhu Fu designed and implemented SEDR. Hang Xu, Huazhu Fu, Kelvin Chong, Mengwei Li, Hong Kai Lee and Jingjing Ling performed data analysis. Hang Xu, Huazhu Fu, Mengwei Li generated figures. Jinmiao Chen, Huazhu Fu, Hang Xu, Kok Siong Ang, Kelvin Chong, Jingjing Ling and Ling Shao drafted the manuscript. Ao Chen and Longqi Liu provided Stereo-seq data. Jinmiao Chen conceptualized and supervised the study.

## Supplementary

**Figure 1.**
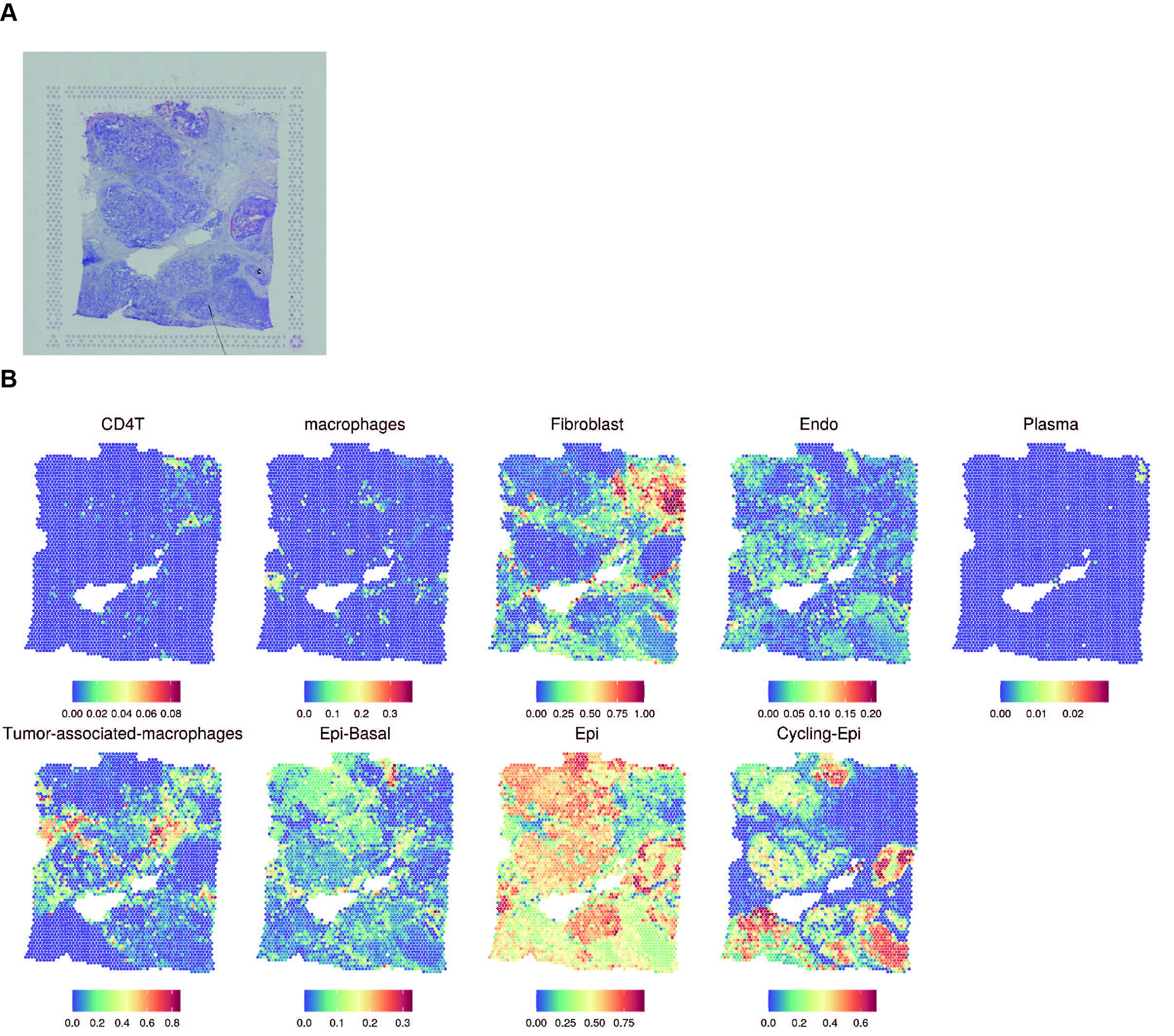
Human breast cancer histology and cell type mixtures of spatial spots. A) H&E staining. B) Seurat3 predicted probabilities of scRNA-seq derived cell types.

**Figure 2.**
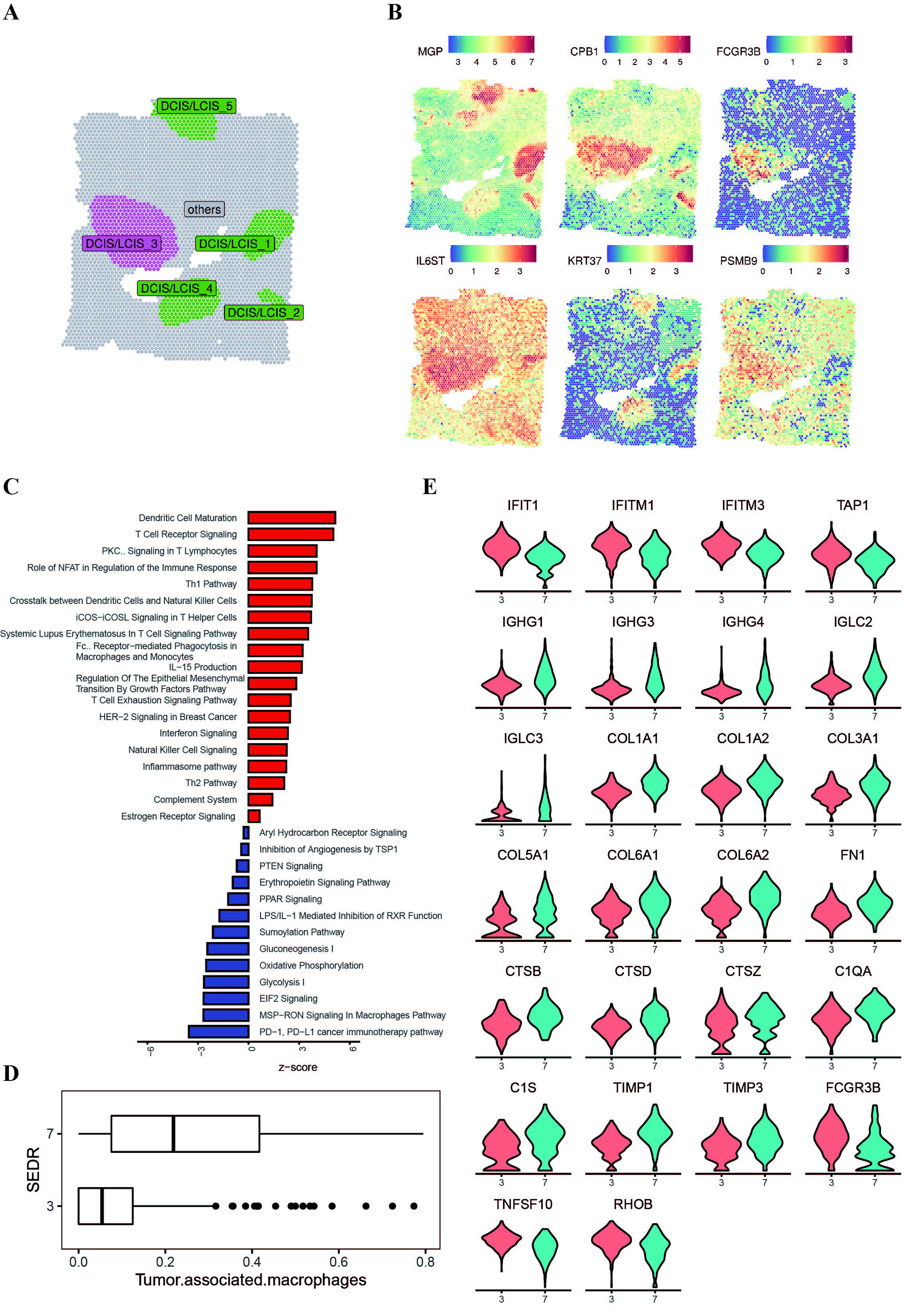
Differentially expressed genes (DEGs) and enriched pathways. A) Locations of DCIS_LCIS_3 and other DCIS_LCIS regions. B) Top DEGs between DCIS_LCIS_3 and other DCIS_LCIS regions. C) Enriched pathways of DEGs between DCIS_LCIS_3 and other DCIS_LCIS regions. Red bars represent pathways up-regulated in DCIS_LCIS_3 D) Percentages of tumor associated macrophages (TAMs) in SEDR cluster 3 (tumor core) and cluster 7 (tumor edge). E) Violin plots of selected DEGs between SEDR clusters 3 and 7.

